# Evaluating risk-taking behaviors in Lycosidae: are the nutritional benefits of sugar water worth the risk of predation?

**DOI:** 10.1101/2022.04.21.489068

**Authors:** Lauren Woyak, Elizabeth McCullagh, Will Wiggins

## Abstract

The risk-taking tendencies and addiction in small vertebrates are often used to understand the processes of larger vertebrates, such as humans. We set out to find whether certain risk-taking behaviors characteristic of addiction remain consistent between vertebrates and invertebrates. Different animals will gorge on sugar water, and addiction-like tendencies are often associated with the overconsumption of sucrose. The goal of my study was to evaluate those risk-taking behaviors and compare invertebrate risk-taking behaviors to vertebrate risk-taking behaviors. The risk-taking behavior we observed was the process of relieving cravings while the risk of predation was present. Through this study we document evidence of wolf spiders feeding on a 25% sucrose solution and explain the wolf spiders’ preference to a sugar water solution over filtered tap water. To evaluate if wolf spiders exhibit addiction like behaviors associated with sucrose, we placed a green-colored, sugar water-soaked cotton ball on the silk of a large, predatory *Tigrosa helluo* wolf spider and a regular, filtered water-soaked cotton ball on the “safe” side of a plastic trial arena and then evaluated the behaviors of the prey spiders. We used a mixture of *Pardosa milvina* and *Rabidosa rabida* as test spiders, with one group given a sugar-water treatment beforehand and the other group was given a green colored control cotton ball with filtered tap water (n=14 for both the sugar water treatment and control groups). The wolf spiders who were predisposed to the sugar water treatment were more likely to cross through their predator’s silk in order to gain access to the sugar water cotton ball than their counterparts who were not given a sugar water treatment prior. These results suggest the risk for predation of the wolf spider is not worth the benefits of the sugar water, unless a wolf spider has been allowed unlimited access to sugar water in advance. When a wolf spider who was allowed unlimited access to sugar water beforehand is presented with a risky situation, they will increase their risk-taking behaviors via crossing through a predator’s silk in order indulge in the sugar water. These results suggest the addictive properties of sugar spans across vertebrates and invertebrates, and potentially opens the door for a cost-efficient and ethically acceptable invertebrate model-organism for examining the risk-taking behaviors associated with addiction.

## Introduction

Food can be used as a natural reinforcer (Hildebrandt and Greif 2013), and the addictive properties of food have been long debated and tested in laboratory studies. Overconsumption of sugars and fats leads to addictive-like behaviors including withdrawal and cravings, as well as an increase in risk-taking (Hildebrandt and Greif, 2013). In fact, when rats over consume sucrose, they are more willing to cross through an electric shock grid to obtain more sucrose and will work harder than the control rats to gain access to foods associated with sucrose (Oswald et al. 2011). However, whether food has addictive properties has been debated and the food addiction model is often criticized to suggest that animals use a more complex mechanism for regulating their feeding habits specifically, there are more interactions than just the neurobiological basis of behavior and addictive behaviors (Hildebrandt and Greif 2013). For example, when an animal is under physical stress such as poor nutritional conditions, they are more likely to engage in risk taking behaviors to meet their nutritional needs (Moran et al. 2020). This further indicates that animals are able to assess their fitness and make decisions based on an internal cost/benefit analysis. Due to the extensive research on spiders using sugar water as a natural reward (Liedtke and Schneider 2013; Wiggins et al. 2018) and changing their behaviors to benefit their fitness (Moran et al. 2020; Persons et al. 2001; Samu et al. 2003; Toft and Wise 1999), they are an excellent model organism for studying behavioral ecology.

Simulated nectar can be made by mixing table sugar with water. When sugar water has the same concentration as natural nectar it can be a useful natural food reward indicating spider preference, and has been long used in studies related to other spider species; particularly jumping spiders (Liedtke and Schneider 2013; Wiggins et al. 2018). Additionally, wandering spiders can enhance their diet and survival by feeding on nectar (Lietzenmayer and Wagner 2017). Jackson et al. (2001) suggested that feeding on nectar may be an overlooked method of feeding in spiders, but it has substantial benefits with low predation risks and low energetic costs.

Lycosidae, more commonly known as wolf spiders, are a wandering spider family known to forage in their environment to find prey (Samu et al. 2002). They are also known to change their foraging methods based on the analysis of the cost/benefit ratio in different environmental conditions and will change their behaviors in the presence of predators (Samu et al. 2002; Barnes et al. 2001). Samu et al. also found wolf spiders will use a mixture of sit-and-wait and active foraging strategies based on prey-density and environmental conditions (2002). Foraging is a metabolically costly activity and because of this, wolf spiders have become great evaluators of a cost/risk to benefit ratio.

Within the Lycosidae family there is a large range in body size, and large wolf spiders often prey on smaller wolf spiders. For example, *Tigrosa helluo* are large (Female: 18-21 mm, Male: 10-12 mm) predatory silk laying wolf spiders who will frequently feed on smaller wolf spiders (Hollenbeck and Elliott 2003; Persons et al. 1999). *Pardosa milvina* are smaller than *T. helluo* with the largest female measuring about 6.2 mm and the largest male about 4.7 mm (Kaston 1981). Developing *Rabidosa rabida* can also be preyed on by *T. helluo* with females averaging 16-21 mm in length and males averaging 13 mm in length (Comstock and Gertsch 1965, Milne and Milne 1980). Since the small prey spiders often live in the same environment as the large predatory spiders, Barnes et al. found prey wolf spiders will decrease their movements in the presence of chemical cues from a larger, predatory wolf spider (2001). A typical chemical cue wolf spiders recognize is the silk dragline from predatory wolf spiders (Barnes et al. 2001).

Based on the addictive nature of sugar for other organisms, nutritional benefit of carbohydrates to spiders, wolf spiders’ ability to analyze cost/benefit scenarios, and the predation of small wolf spiders by larger wolf spiders, we decided to evaluate whether or not the benefits of feeding on sugar water were greater than the risk of passing through a predator’s silk. The detection of predatory silk, analysis of what the predatory spider ate, and perceived predation risks are all considerations before making a decision (Persons et al. 1999). The aim of my study was to determine if wolf spiders had an affinity for sugar water, like other wandering species previously tested (Lietzenmayer and Wagner 2017), and to determine if *P. milvina* and *R. rabida* would risk potential predation for the nutritional boon of sugar water. We predicted the wolf spiders would choose the sugar water even with the potential to be preyed upon. We hypothesized that since sugar water is a high-source of nutrients for a spider, the small, prey spiders (*P. milvina* and *R. rabida*) who learned to associate a green colored cotton ball with sugar water would be more willing to pass through a predator’s (*T. helluo*) silk and the control spiders (given no sugar water) would not pass through the silk.

## Methods

### Spider collection and care

We collected spiders from the Lycosidae family by using a spotlighting technique while walking through Sanborn Lake Park in Stillwater, OK. The spotlighting technique illuminates the eyes of the wolf spider and allows for easy identification of their location in the dark. The spiders were collected in the fall of 2020 and the spring of 2021. The spiders were kept in a room (about 22°C) and had a light cycle of 14:10 (light: dark).

Each spider was kept in an its own individual round, opaque and plastic container measuring 8 cm tall with a diameter of 11 cm. All spiders were kept in close proximity, potentially exposing the spiders to mechanical and chemical cues from the other spiders (Uetz and Roberts 2002). Spiders were placed in three groups: predatory spiders (*T. helluo*), control spiders, and experimental spiders both of which contained a mix of *P. milvina* and *R. rabida*. The predatory spiders were about four to five times larger than the control and experimental spiders. All spiders were given size appropriate crickets, *Gryllodes sigillatus*, (2-3 crickets twice a week) and colorless cotton balls with filtered tap water. However, one week before the experiments, the *T. helluo* were starved in order to ensure there would not be prey-type chemical cues present in their silk draglines. The experimental spiders additionally had a cotton ball with green colored sugar water. The sugar water was a 25% sucrose solution, which is typical for a flower with nectar (Chalcoff et al. 2005; Wolff 2006). The sugar solution was dyed green with eight drops of McCormick green food coloring to induce association of the sucrose solution with a green color in the experimental group. A green color was used because wolf spider eyes are sensitive to blue-green wavelengths (DeVoe et al. 1969). The cotton ball with sugar water was removed from the container when the spiders were feeding on crickets, to ensure the crickets were not feeding on the sugar water. The control spiders were also given green colored cotton ball, except we used filtered tap water also dyed with eight drops of McCormick green food coloring. The green cotton ball with filtered tap water is a control for color or food dye preference of wolf spiders when choosing a cotton ball to drink/feed from.

### Testing Lycosidae affinity for sucrose solution

Before the experimental spiders were given a consistent supply of sucrose solution, we first tested to see if the spiders would drink the sucrose solution. They were given two colorless cotton balls, one with the sugar water and one with filtered tap water. The spiders were allowed to acclimate for five minutes, and then observed for 15-16 minutes. We then recorded the average percentage of time the spiders spent on each cotton ball for day one and day two, and analyzed the difference in the means using a t-test (alpha = 0.05). After one day, spiders continued to show preference in drinking from the sugar water cotton ball (Fig. 2).

### Risk-taking behaviors

Wolf spiders rely on chemical cues from silk and excretions to detect the presence of predators (Barnes et al. 2001), so we observed the risk-taking behavior of crossing through a predator’s silk to retrieve sucrose solution. All trials were conducted in a round, clear plastic container with a glass lid. The container was 9 cm tall and had a diameter of 21 cm. The container was divided into two equal halves by a cardboard divider. The cardboard divider had petroleum jelly on it to discourage climbing. A large predatory spider was placed in one half of the container and allowed to walk around, and lay silk, for 10 minutes. After the 10 minutes, the large predatory spider was placed back in its respective container, and the center divider was removed from the trial arena. Once the center divider was removed, a colorless cotton ball with filtered tap water was placed on the side of the trial arena without the predatory spider silk, and the green cotton ball with sucrose solution was placed on the side of the arena where the large predatory spider had just laid silk. To rule out directional bias, the cotton balls were alternated between North and South with the green cotton ball starting in the North position and in the next trial it would be in the South position. The small prey spider was then released into the center of the trial arena and allowed to acclimate for 5 minutes. After the acclimation period, the spider was recorded for 10 to 20 minutes to observe its time spent on the sugar water and silk side of the arena, and how much time was spent on the filtered tap water side of the arena (Fig. 1). Trials were completed with both the experimental group and the control group (n=14 for each group).

**Fig. 1.**
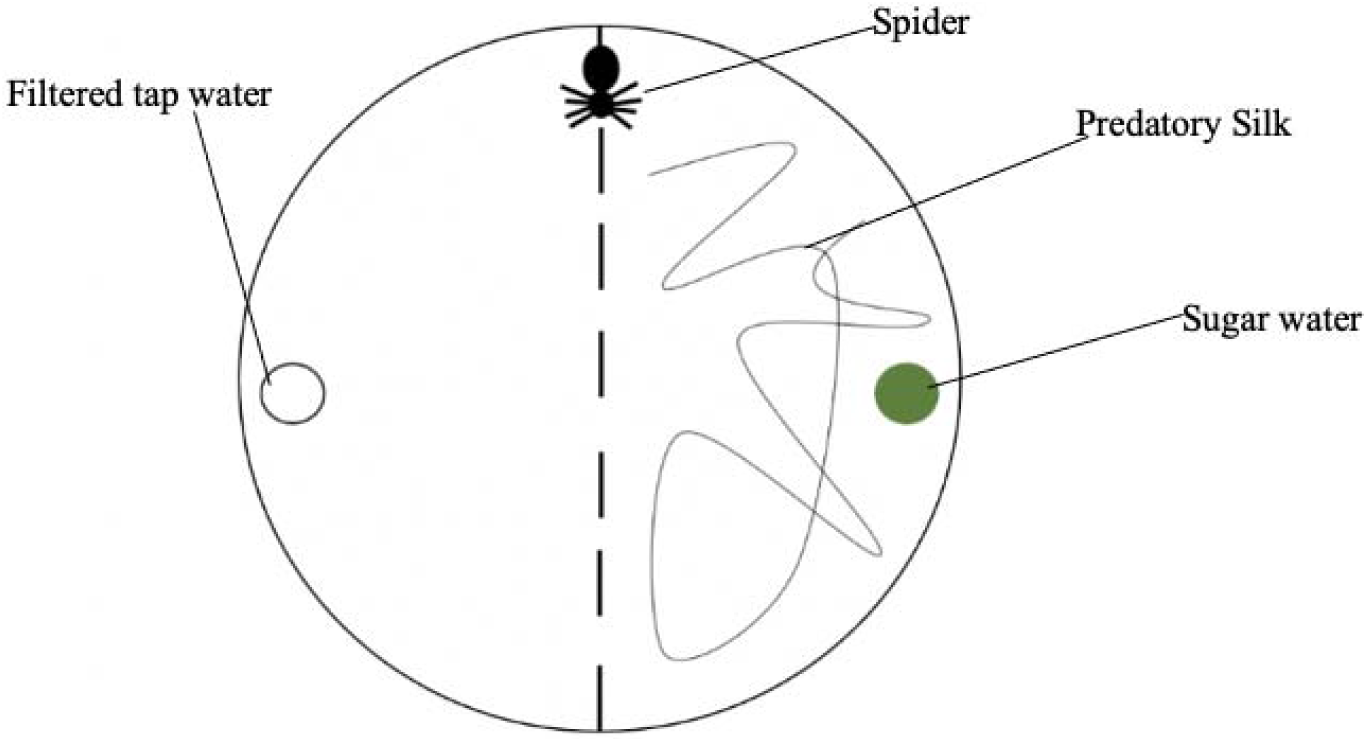
Schematic drawing of trial arena with prey spider. When the prey spider was placed into the arena, the center divider was removed, but the center of the arena was still marked. The direction the sugar water was placed in (North vs. South) was alternated, but ultimately determined to be statistically insignificant (p=0.9412).

After each trial, we removed the spider and cleaned the container with 91% isopropyl alcohol to remove all the silk and scent from the previous spiders. Trials were conducted in both the fall of 2020 and the spring of 2021. The same procedure was used for both sets of trials. Videos of the spider movements were analyzed to determine if spiders given sugar water prior the trial chose to spend more time on the side of the arena that had the predatory silk and sugar cotton ball compared to spiders only given filtered tap water. The data from the first set of trials in fall of 2020 was analyzed with a t-test to determine if the mean time spent on the sugar side of the trial arena was significantly different from the mean time spent on the sugar side of the arena between the control and experimental groups. After data was collected for the second set of trials in the spring of 2021, the same analysis was used to again determine if the means between the control and experimental groups significantly differed.

## Results

To author’s knowledge, no articles have indicated that wolf spiders will drink sugar water nor have a preference to sugar water over filtered water. Before risk-taking trials were performed, we evaluated whether or not the Lycosidae had a preference for sugar water over regular filtered water. After being observed for 15-16 minutes both on day one and day two, the wolf spiders showed a statistically significant preference for sugar water with a t-value of 26.624, a t-critical two-tail value of 12.706 and a p-value of 0.0238 (n=8, alpha=0.05) (Fig. 2). Spiders were then evaluated for risk-taking behaviors within two weeks after the preference trials. A t-test was conducted to determine the significance in the difference of means between the experimental group and the control group (Figs. 3,4). The spiders who were priorly exposed to sugar water were more likely to cross through the predator’s silk than their counterparts who had not been previously exposed to the sugar water treatment in both the preliminary trials in fall 2020, and the second set of trials in spring 2021. The preliminary trials had a t-score of -2.46 with a t-critical two-tail value of 2.45, and p=0.0488 (n=7 and n=6 for control and experimental groups, respectively). The second set of trials had a t-score of 14.75 with a t-critical two-tail value of 12.71, and p= 0.04310 (n=7 and n=8 for control and experimental groups, respectively). Both t-tests used alpha=0.05. For both trials, the mean percentage of time spent on the sugar water side for the experimental groups was around 60%: 68.04% for trial 1 group and 61.26% for trial 2 group (Figs. 3,4). The mean percentage of time spent on the sugar water side for the control group was about 35%: 37.24% for trial 1 group and 34.38% for trial 2 group (Figs. 3,4). Using JMP (SAS Inc. 2016), a chi-square test was used to determine if the direction in which the sugar water was placed (North vs. South) influenced the outcome. It was not significant with a chi-square value of 0.005 and p=0.9412 (n=7 and n=8 for control and experimental groups, respectively).

**Fig. 2.**
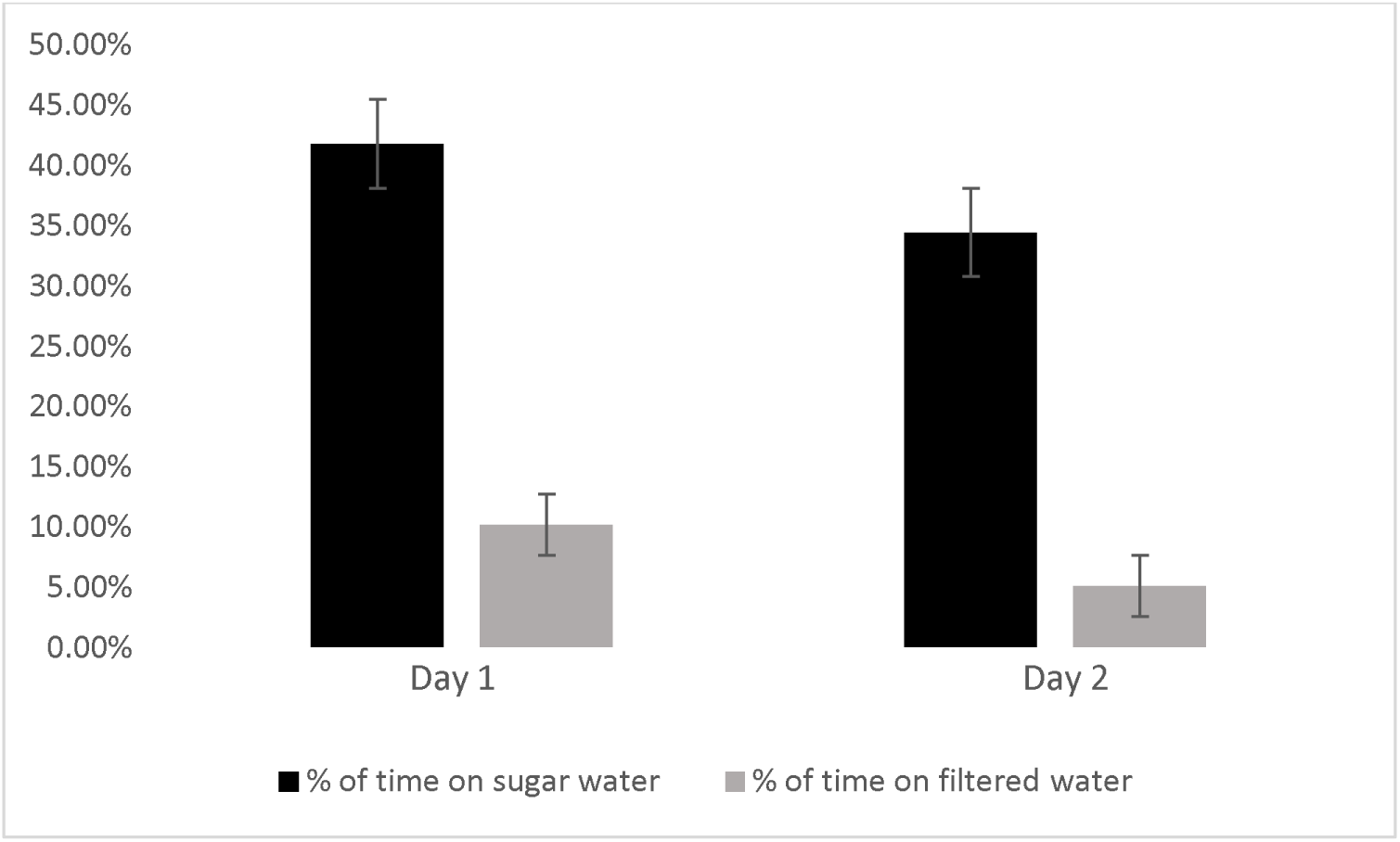
Mean percentage of time the wolf spiders spent on either the cotton ball with sugar water or the cotton ball with filtered tap water on days one and two. The spiders were given five minutes to acclimate and were then observed for 10-20 minutes. The spiders spent less time on either cotton ball on day two, but the difference between the means both days was still significant, showing preference for the sugar water (t-stat= 26.624, t-critical two tail= 12.706, p=0.0238).

**Fig. 3.**
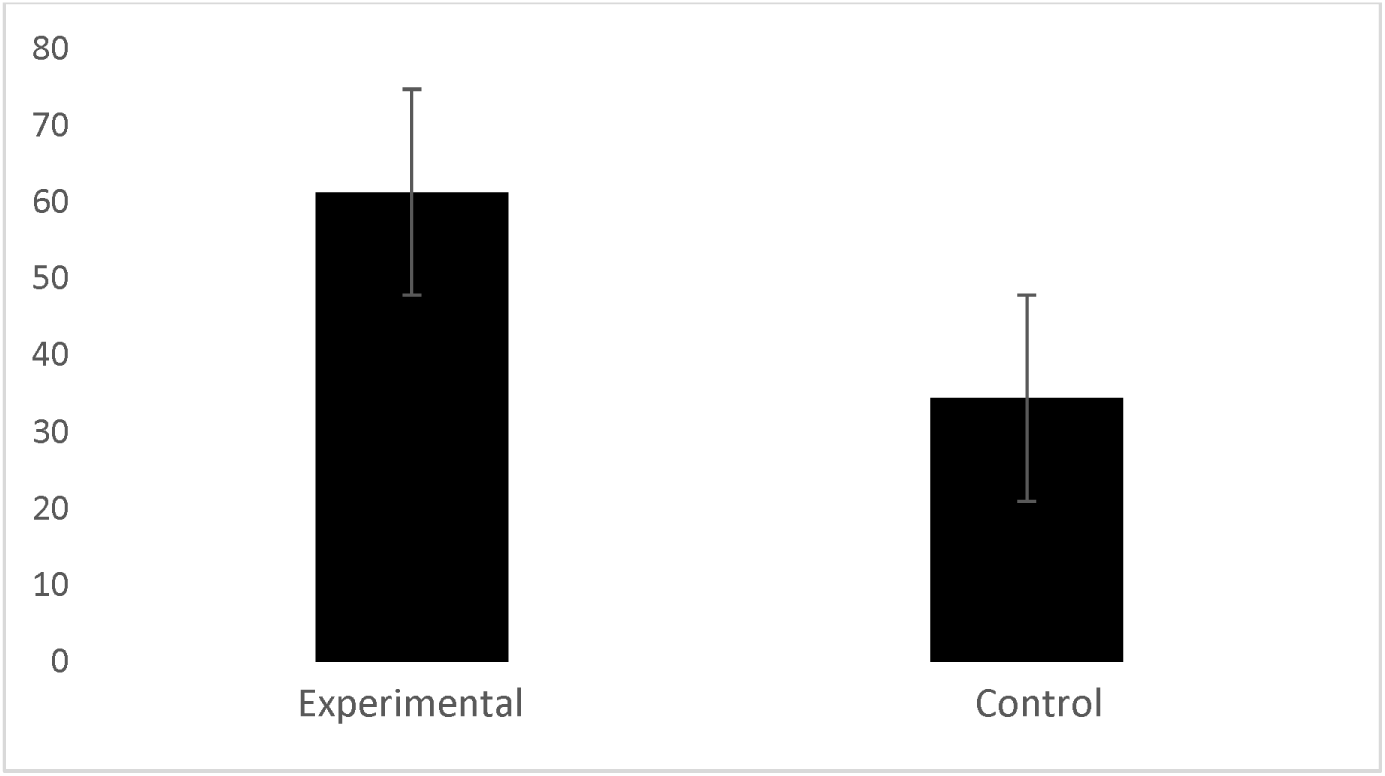
T-test results for mean % of time spent on sugar side of arena for experimental group vs. control group in fall of 2020 data group. The spiders who were in the experimental group and given a sugar treatment prior to testing spent more time on average on the sugar side of the trial arena in comparison to the control group (p=0.0488).

**Fig. 4.**
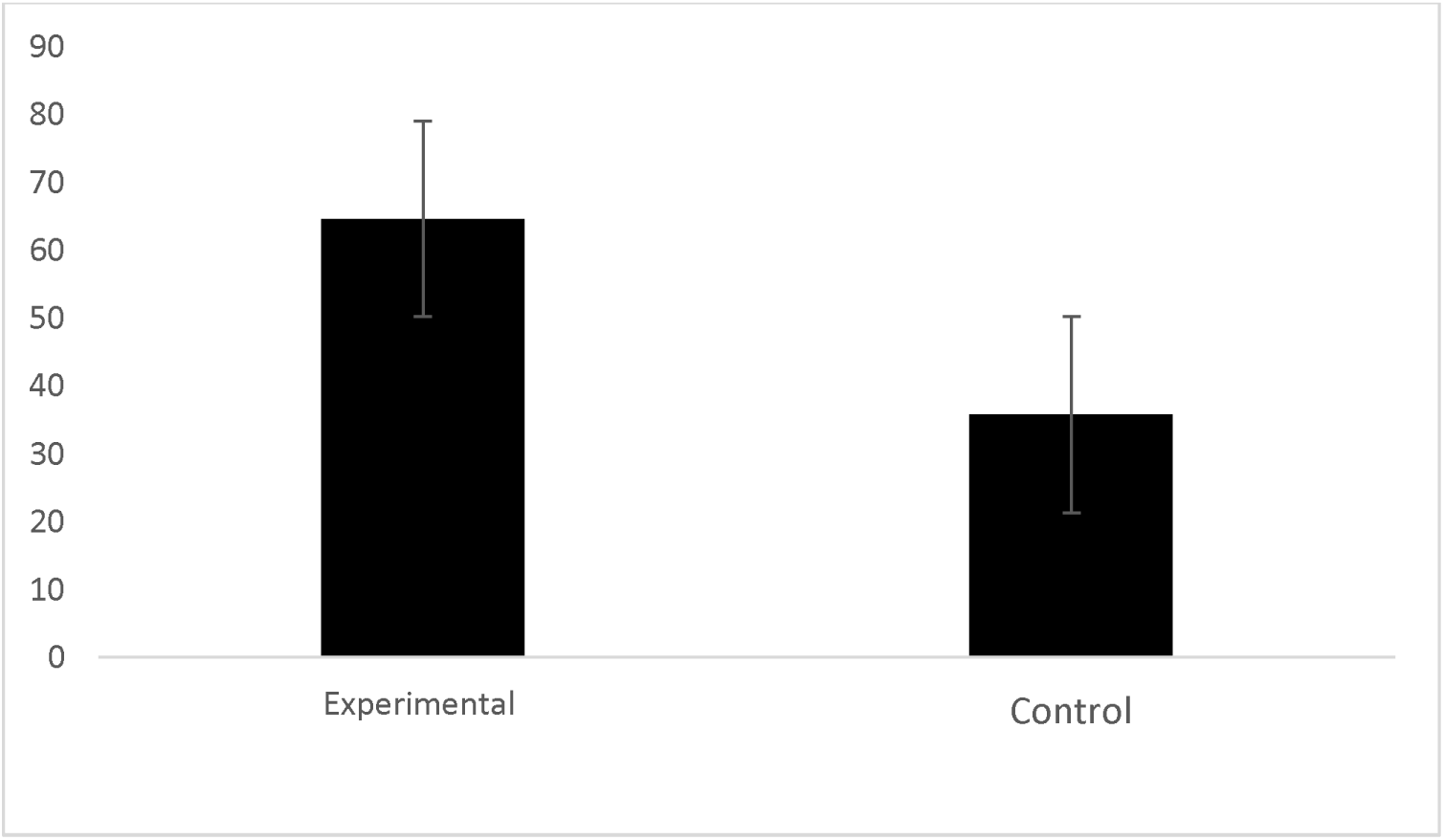
T-test results for mean % of time spent on sugar side of arena for experimental group vs. control group in spring of 2021 data group. Again, the spiders who were given a sugar water treatment prior, spent more time on average on the sugar side of the trial arena than the spiders who were not given a sugar water treatment prior (p= 0.04310).

## Discussion

Sugar water has been used as a natural reward (Liedtke and Schneider 2013) and demonstrated increased survival in jumping spiders (Wiggins unpublished). However, these relationships have not been explored in wolf spiders, and this is the first paper to indicate addictive like behaviors in arachnids. The wolf spiders showed a consistent preference for the sugar water after 24 hours (p=0.0238) (Fig. 2). This is consistent with data from Jackson et al., which indicates jumping spiders will use sugar water as a natural food reward (2001), and in correspondence with my data, suggests that the use of sugar water as a natural food reward is applicable to the wolf spider species.

The purpose of this study was to observe the risk-taking behaviors of Lycosidae after they were exposed to a sugar water treatment, and then compare those behaviors to the Lycosidae which were not exposed to a sugar water treatment. We predicted the wolf spiders would indulge in the sugar water, even with predator cues present. Persons et al. (2001) described how *P. milvina* will decrease their movements in the presence of fresh silk laid by *H. helluo*, which the control spiders did. The control spiders chose to be on the side of the arena without freshly lain silk from *H. helluo*. However, the experimental spiders who have associated the green cotton ball with sugar water would exhibit risk-taking behaviors to obtain the sugar water. This supports my hypothesis that the wolf spiders who learned to associate the color green with sugar water would be more likely to risk predation to access the sugar water than the wolf spiders who did not learn to associate the color green with sugar water. Spiders have excellent chemoreceptors and mechanoreceptors to aid them in sensing their environment, and all results indicate they were aware of the predator silk inside the arena (Uetz and Roberts 2002).

This awareness and willingness to undergo a risky situation for a substance is in line with our understanding of addiction behaviors, where individuals will increase risk-taking behaviors to obtain their addictive substance (Hildebrandt and Greif 2013; Moran et al. 2020; Oswald et al. 2011, Voon et al. 2014). These results are consistent with Oswald et al.’s findings which explained when rats overconsume sucrose, they were more likely to cross through an electric shock to receive the sucrose than their control counterparts who did not overconsume sucrose (2011). From this, we conclude that the addictive properties of sugar can affect vertebrates and invertebrates alike.

This is an exciting find, and may help us better understand addiction behavior, anatomy, and physiology. Using spiders as a model species allows us an opportunity to explore the similarities and differences of addiction between vastly different organisms. In future studies we would be interested in examining if addicted spiders will exhibit risky behaviors for green cotton balls filled only with filtered tap water and explore the neurochemical changes that occur in a spider’s brain when addicted to sugar. Avena et al. demonstrated how rat brains during a sugar dependency are similar when viewing the rat’s brain during drug addiction (2007).

It is important to understand this connection in order to explain how using sugar as an addictive substance can be sufficient to evaluate certain risk-taking behaviors that occur with addiction. Because the addictive properties of sugar seem to be consistent throughout species and families in the animal kingdom, wolf spiders may become a convenient, and cost efficient, model organism for studying basic risk-taking tendencies often seen in relation to addiction or eating disorders.

## Acknowledgements

I would like to thank all of my friends who were always around to help me collect spiders late at night, listen to my progress, and be additional eyes for all of my drafts. I could not have done it without you all, LW.

